# CASCADE recovers promoter-associated regulatory motifs from cell-type-resolved DNA language-model attributions

**DOI:** 10.64898/2026.07.16.738979

**Authors:** Ali Farghadan, Robert J. Schmitz, Scott A. Jackson, Ethan Pickering

## Abstract

Gene expression is governed by regulatory DNA and their associated trans factors acting in specific cell types, yet the sequences underlying this control remain poorly mapped in plants. Genome-pretrained DNA language models provide a route to interrogate regulatory sequence directly, but their attributions have largely been interpreted using bulk or whole-tissue data, and standard attribution pipelines can preferentially highlight sequences downstream of the transcription start (TSS) site rather than promoter-associated signals. Here, we train a celltype-resolved sequence-to-expression model from a single-cell soybean (*Glycine max*) atlas by coupling a soybean-adapted Genomic Pre-trained Network (GPN) to a shared sequence encoder with 66 cell-type-specific output heads. Across 38,339 protein-coding genes, the model achieves a mean per-cell-type, across-gene Pearson correlation of 0.683 and, recast as a highversus-low expression classification, reaches an area under the ROC curve of 0.92 to 0.97 across tissues, at or above dedicated plant sequence models. We then introduce Context-Aware Significance of Cross-gene Attribution for Discovering Elements (CASCADE), a position-specific statistical framework for identifying model-derived candidate regulatory elements from *in silico* saturation mutagenesis. Relative to the pooled null used by TF-MoDISco, CASCADE shifts motif recovery from downstream of the transcription start site toward promoter sequence, with 77% of CASCADE-exclusive motifs, compared with 12% of TF-MoDISco-exclusive motifs, falling within the promoter. Applied across the atlas, CASCADE identifies approximately 1.39 million candidate elements spanning broadly active, tissue-restricted and cell-type-restricted classes. Together, these analyses establish a position-aware approach for extracting promoter-associated regulatory hypotheses from sequence models and generate a cell-type-resolved map of candidate *cis*-regulatory elements.

## 1 Introduction

Predicting gene expression from DNA sequence has matured into a quantitative tool for regulatory genomics. The field has advanced from Enformer (Avsec et al., 2021a) to Borzoi (Linder et al., 2025) and AlphaGenome (Avsec et al., 2026) as receptive fields and parameter counts have grown. As these models have become sufficiently accurate, the question of interest has shifted from how well they predict to what they have learned. We now ask which bases in a gene’s neighborhood carry information about its expression, and whether that information is shared across all cell types or specific to particular cell types. A sequence-to-expression (S2E) model is, after training, a fixed differentiable function from DNA to expression, and perturbing its input reveals which positions the prediction depends on. This makes attribution a route to a sequence-resolved account of regulation, complementary to the experimental maps of accessible chromatin and transcription-factor occupancy on which it can be evaluated.

However, attribution magnitude is strongly position dependent around genes. Regions downstream of the transcriptional start site often produce larger perturbation effects than upstream promoter sequence, so that a position-agnostic significance threshold can preferentially identify downstream features even when the biological question concerns promoter *cis*-regulation (Mendoza-Revilla et al., 2024; Peleke et al., 2024; Michielsen et al., 2024). Recovering promoter-associated grammar therefore requires comparing attribution at each position against an appropriate position-matched background. In this study, *promoter* refers operationally to the 2,000 bp immediately upstream of the TSS and *gene body* to the 2,000 bp immediately downstream, the two halves of the fixed 4,000-bp window we analyze around each gene. These are positional windows relative to the TSS, not annotated promoter or gene-body boundaries.

*In silico* saturation mutagenesis (ISM) and related attribution methods have been used to recover regulatory grammar directly from trained models. Promoter-focused analyses of human models recover the core-promoter elements, including the TATA box. The positional effect can be large in a single sequence background and more modest genome wide (Karollus et al., 2023). The masked-language model GPN (Genomic Pre-trained Network) (Benegas et al., 2023) is trained on *Arabidopsis* and related Brassicales without any expression labels. It recovers start and stop codons and roughly 160 promoter motifs as position-locked signals. In soybean and other crops, a bulk model of transcriptional initiation recovers 27 core promoter motifs, including the TATA box and the Initiator, by *in silico* mutagenesis (Gao et al., 2025). Coverage models that predict RNAseq from sequence additionally learn splicing and polyadenylation signals (Linder et al., 2025). These results establish that trained sequence models capture signals associated with established regulatory and structural sequence features. They are, however, drawn almost entirely from bulk or aggregated data and from mammalian or whole-tissue settings. Although sequence models learn genuine signals in untranslated and gene body regions, these strong positional signals can dominate attribution-based motif discovery and obscure signal from promoter-associated regulatory grammar. It remains unclear how to distinguish promoter *cis*-regulatory sequence from coding sequence attribution, particularly when regulatory effects must also be resolved across cell types.

The difficulty is sharpest precisely in the cell-type-specific regime that single-cell data expose. Aggregate accuracy metrics are dominated by broadly expressed genes. Stratified analyses show that current sequence models predict the conditional, cell-type-defining component of expression poorly (Karollus et al., 2023). Cell-state-conditioned decoders trained on millions of single cells only partly address this limitation (Hingerl et al., 2025; Lal et al., 2026). Models that predict cellpopulationor cell-type-specific expression from sequence most nearly address this, and some interpret that sequence by *in silico* mutagenesis or variant scoring (Michielsen et al., 2024; Zhou et al., 2025), yet none maps, one cell type at a time, which promoter-proximal sequence a model uses across a multi-cell-type atlas. A single number therefore conveys little about *where* a model is informative. What is missing is a systematic account of promoter-proximal sequence attribution resolved across the cell types of one organism. It should report which positions a model uses and how broadly each acts.

Soybean makes these questions both consequential and informative. It is a leading global source of plant protein and oil and fixes atmospheric nitrogen through a symbiotic root nodule (Roy et al., 2020), so the regulatory programs that distinguish its organs directly influence agronomic traits (RodrÍguez-Leal et al., 2017). Its genome has undergone two whole-genome duplications,leaving roughly three-quarters of genes in duplicate (Schmutz et al., 2010), and a major route by which such duplicates diverge is change in their *cis*-regulatory sequence rather than their coding sequence (Fang et al., 2023; Li et al., 2025), which a cell-resolved map of candidate regulatory elements may help expose. More broadly, much of the trait and domestication variation in crops is non-coding (Meyer and Purugganan, 2013), so an element atlas paired with a sequence model that scores variant effects offers a route to prioritize candidate regulatory variants and predict the cell types in which their effects may occur.

Here, we develop a position-aware framework for recovering promoter-associated regulatory sequence from a cell-type-resolved sequence-to-expression model. We couple a soybean-adapted Genomic Pre-trained Network to a shared sequence encoder with 66 cell-type-specific output heads and use *in silico* saturation mutagenesis to quantify the predicted effect of sequence perturbations around 38,339 genes. We then introduce CASCADE, which evaluates attribution against a cross-gene, position-specific null rather than the pooled null used by TF-MoDISco (Shrikumar et al., 2018). This shifts motif recovery from coding-proximal sequence toward upstream promoter regions, while preserving established motif recovery on a human core-promoter benchmark. Applied across the soybean atlas, CASCADE identifies candidate elements ranging from broadly active to tissue- and cell-type restricted, many not found with TF-MoDISco, providing a cell-type-resolved map of promoter-associated regulatory hypotheses in a major crop.

## 2 Results

### 2.1 CASCADE recovers promoter-associated regulatory sequence from positionally biased attribution scores

Attribution-based motif discovery is strongly influenced by position around the transcription start site, and a position-specific cross-gene null shifts candidate recovery from gene body sequence into the promoter. To identify promoter-associated sequences while accounting for this positional structure, we developed Context-Aware Significance of Cross-gene Attribution for Discovering Elements (CASCADE). CASCADE evaluates attribution scores at each position against the distribution observed at the same TSS-relative position across genes, rather than against a single pooled background. When applied to soybean sequence-to-expression attributions, this position-specific comparison shifts the seqlets it identifies, contiguous DNA intervals selected from an attribution track due to their aggregate attribution score, away from the region immediately downstream of the transcriptional start site and toward upstream promoter sequences (Figure 1).

**Figure 1:**
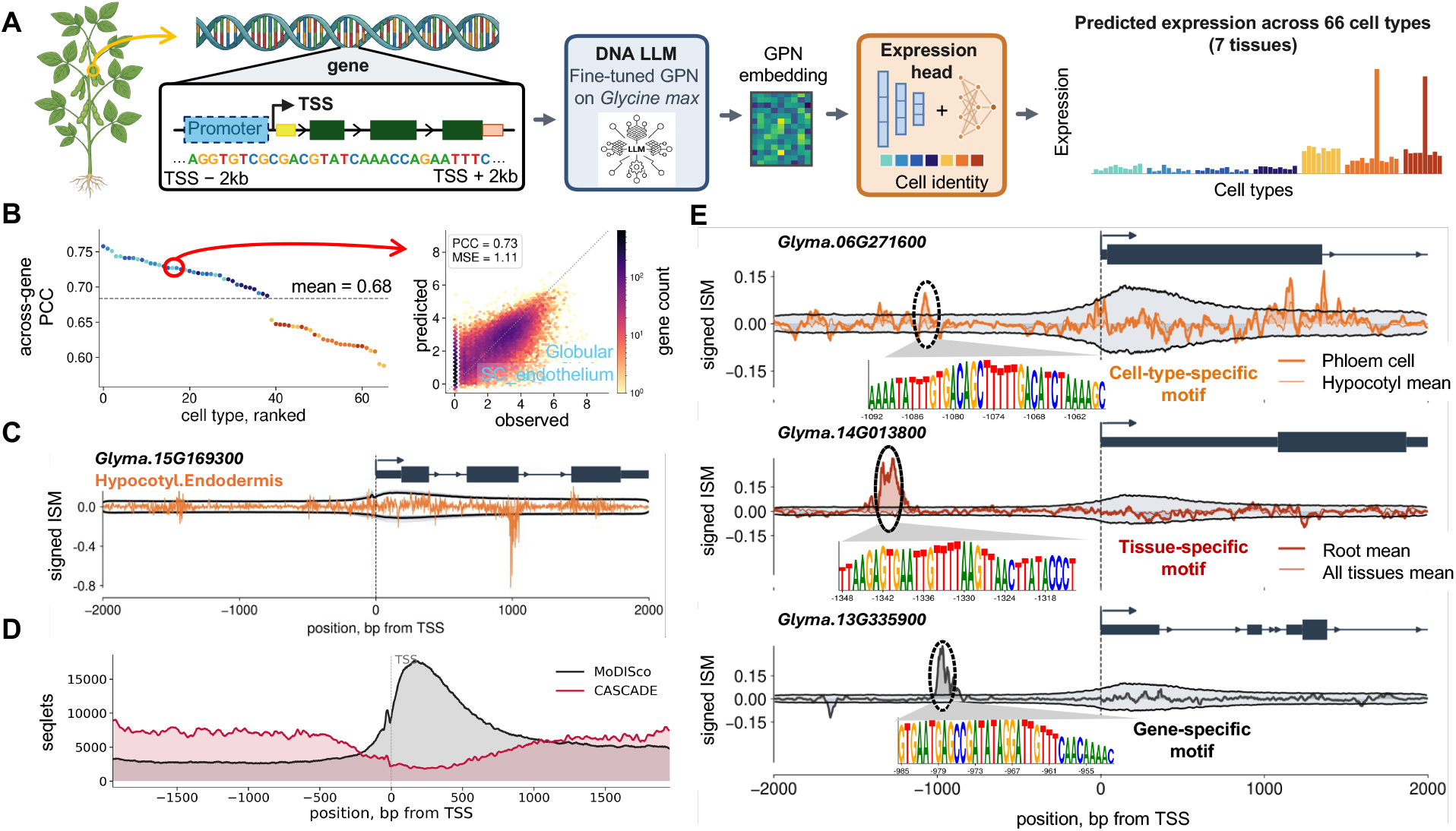
CASCADE converts cell-type-resolved sequence attributions into promoterassociated candidate regulatory elements across multiple cellular contexts. **A** Soybeanadapted GPN embeds the 4-kb genomic window centered on each gene’s TSS at nucleotide resolution, and a shared convolutional encoder with 66 cell-type-specific output heads predicts expression across the atlas. *In silico* saturation mutagenesis perturbs each position within this window to generate a signed attribution track for every gene and cell type. **B** Per-cell-type, across-gene test Pearson correlations under chromosome-blocked 10-fold gene-level cross-validation, with one point per cell type colored by tissue and the mean shown by the dashed line; right, predicted versus observed log_10_ *p*(CPM) for a cell type at the 75th percentile of predictive performance. **C** Example signed-*in silico* saturation mutagenesis attribution track for one gene and cell type, shown relative to the cross-gene distribution at each TSS-relative position. **D** Positional distribution of seqlets under the pooled TF-MoDISco null and the position-specific CASCADE null. Although both operate on the same attributions, the pooled null concentrates seqlets immediately downstream of the TSS, whereas CASCADE shifts them toward upstream promoter sequence. **E** Cellular-context resolution of CASCADE detections. Applying the position-specific null independently to each celltype attribution track identifies elements detected broadly across cell types, restricted to one tissue or confined to one or two cell types. Example loci show the signed-*in silico* saturation mutagenesis track, position-specific significance threshold, candidate element and local sequence motif.

The attribution tracks supplied to CASCADE were generated from a cell-type-resolved sequenceto-expression model trained on a single-cell soybean atlas (Zhang et al., 2025). A soybean-adapted GPN (Benegas et al., 2023) provides nucleotide-resolution embeddings of the 4-kb window centered on each gene’s TSS, which are processed by a shared convolutional encoder with 66 cell-type-specific output heads (Figure 1 A). Across 38,339 protein-coding genes, the model achieves a mean per-cell-type, across-gene Pearson correlation of 0.683 under chromosome-blocked cross-validation (Figure 1 B). Current plant sequence models are typically evaluated as binary high-versus-low expression classifiers (Peleke et al., 2024). When evaluated under this setting, the same held-out predictions separate high- from low-expressed genes in every tissue at an area under the ROC curve of 0.92 to 0.97, at or above the reported range for such models (Figure S1). The ultimate goal, however, is to predict the exact expression level rather than a bin, which would assist concentration-aware regulatory network inference. *In silico* saturation mutagenesis then produces a signed attribution track for every gene and cell type, quantifying the predicted expression change associated with perturbation of each position (Figure 1 C). These cell-type-resolved attribution tracks provide the input to CASCADE rather than constituting the primary result themselves.

CASCADE extends this positional comparison across cell types, allowing the same candidate sequence to be resolved according to the cellular contexts in which it is detected. The method is applied independently to the attribution track of each cell type, producing a set of seqlets. Overlapping intervals are then merged across cell types, and their occupancy patterns distinguish sequence elements detected broadly across the atlas from those restricted to a tissue or to one or two cell types (Figure 1 E). This framework therefore combines two forms of context: TSS-relative position determines the appropriate cross-gene attribution background, whereas celltype-resolved model outputs determine the biological contexts in which each candidate element is identified.

### 2.2 A position-specific null reallocates motif discovery from downstream of TSS to the promoter upstream

Holding the attributions and clustering procedure fixed, replacing the pooled TF-MoDISco null with the position-specific CASCADE null reallocates motif discovery from coding-proximal sequence toward the upstream promoter. A motif-discovery pass first identifies significant seqlets and then clusters those seqlets into recurring motifs. To isolate the effect of the null, we used identical cellaveraged in silico saturation mutagenesis attributions, window length, flank, target false-discoveryrate (FDR), and downstream clustering, changing only whether significance was evaluated against a pooled or position-specific background.

The two procedures recover similar total numbers of seqlets but distribute them differently around the TSS. TF-MoDISco returns 354,155 seqlets and CASCADE returns 347,316, indicating that the position-specific null primarily reallocates seqlets rather than simply increasing their total number. Of the resulting motifs, 18 match one-to-one between the procedures at a contributionweight-matrix correlation of at least 0.80 (Figure 2 A). CASCADE returned 42 repressor and 19 activator motifs, compared with 30 repressor and 29 activator motifs under TF-MoDISco. Among the 18 motifs shared between procedures, per-motif seqlet counts remained correlated in rank (Spearman *ρ* = 0.79, *p* = 1 *×* 10^*−*4^, *n* = 18; Figure 2 C), with counts for the eight highest-confidence pairs shown in Figure 2 B, although most points fell below the diagonal, indicating that CASCADE assigned fewer seqlets to these shared motifs while reallocating seqlets to regions upstream of the TSS. Although their sequence logos are highly similar, TF-MoDISco places the contributing seqlets primarily near or downstream of the TSS, whereas CASCADE identifies instances of the same motif predominantly within the upstream promoter.

**Figure 2:**
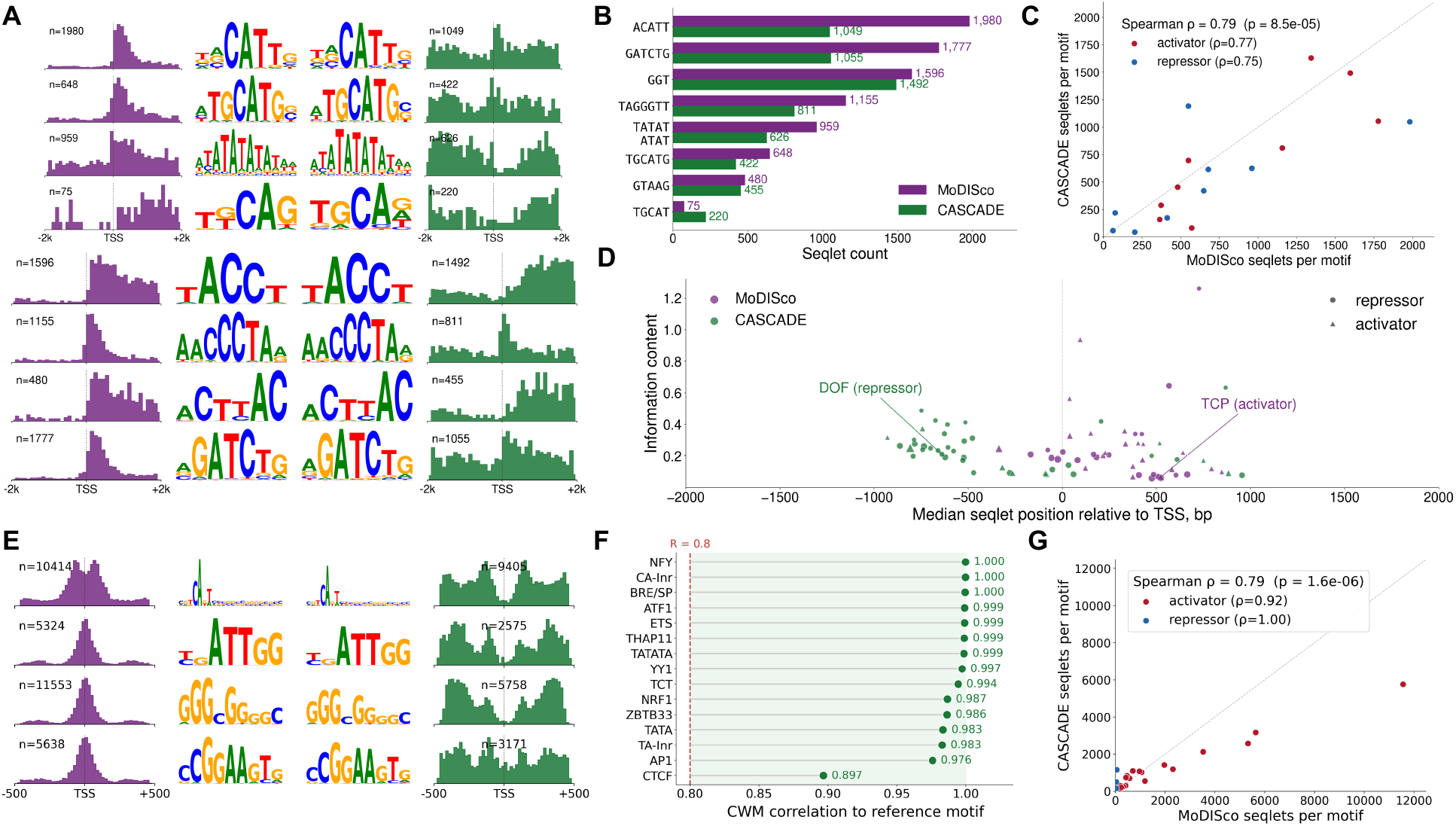
A position-specific null relocates soybean motif discovery from downstream of the TSS into the promoter while preserving established core-promoter motif recovery on a human benchmark. CASCADE replaces the pooled, position-flat null used by TF-MoDISco with a position-specific null of TF-MoDISco with a per-position FDR null, holding window, flank, target FDR and clustering fixed, so differences reflect the null alone. Throughout, MoDISco (or the deposited reference) is purple and CASCADE green. (**A**–**D**) soybean. **A** Shared motifs (one-to-one contribution-weight-matrix match); the two seqlet-position distributions relative to the TSS flank each logo pair, near-identical in shape but placed downstream by MoDISco and in the promoter by CASCADE. Each pair is labeled with its best Tomtom-matched transcription-factor family (q *≤* 0.05) or “unannotated” if no confident match was found. **B** Per-motif seqlet counts for the shared motifs. **C** Rank agreement of seqlet abundance among shared motifs; points below the diagonal indicate motifs to which CASCADE assigns fewer seqlets. **D** Method-exclusive motifs by median seqlet position relative to the TSS and per-position information content (point size, seqlet count; low-complexity motifs excluded), MoDISco-exclusive downstream and CASCADEexclusive upstream; marker shape distinguishes repressor from activator motifs, with one named example of each highlighted. **E**–**G** Human K562 (ProCapNet benchmark). **E** Best-recovered named factors, with seqlet-position distributions relative to the PRO-cap summit flanking each logo pair. **F** Recovery of every annotated core K562 factor, ranked by contribution-weight-matrix correlation to the reference (dashed line, 0.8). **G** Rank agreement of shared-motif seqlet counts; points lie close to the diagonal.

The positional difference is strongest among motifs recovered exclusively by one procedure. The 41 TF-MoDISco-exclusive motifs have a median position of +375 bp relative to the TSS, whereas the 43 CASCADE-exclusive motifs have a median position of *−* 521 bp. Consequently, 77% of CASCADE-exclusive motifs fall in the promoter, compared with 12% of TF-MoDISco-exclusive motifs (Figure 2 D). The two sets have similar per-position information content–0.20 and 0.23 bits, respectively–and CASCADE-exclusive motifs match annotated transcription-factor families under Tomtom at least as frequently as TF-MoDISco-exclusive motifs (31*/*43 versus 26*/*41). Thus, the promoter shift is not explained by a loss of motif complexity or recognizable transcriptionfactor grammar.

Because a position-specific null is more permissive where background attribution is low, part of this promoter shift follows directly from the statistical design. The matched motif quality and transcription-factor-family recovery indicate that CASCADE is recovering structured promoter-associated sequence rather than indiscriminately lowering the significance threshold.

To test whether CASCADE generalizes across species, we applied it to human, where independently annotated core-promoter transcription factors provide a stringent external benchmark (Cochran et al., 2024). On the human benchmark, CASCADE preserves established regulatory grammar while introducing only limited changes relative to the deposited TF-MoDISco results. This benchmark is an external positive control in a well-characterized human system, separate from the soybean analysis: we ran the identical comparison on the humanK562 PROcap model of ProCapNet (Cochran et al., 2024), whose core-promoter transcription factors are annotated. Figure 2 E,F show that CASCADE recovers all 43 reference activator motifs, with 41 of the 43 matching the deposited references at a contribution-weight-matrix correlation of at least 0.8. Among the annotated core transcription factors, recovery is uniformly high, with a median correlation of 0.997 and a minimum of 0.897. That minimum is CTCF, a mammalian insulator factor present in the human benchmark but with no plant counterpart; its long, low-information flanks make it the hardest core factor to match by correlation, and the per-position null slightly fragments it, rather than failing to recover it. For the best-recovered factors, CASCADE’s seqlets nonetheless shift away from the PRO-cap summit itself and into the immediately flanking sequence (Figure 2 E): these factors carry strong, near-universal initiator attribution at the summit in almost every promoter, so the per-position null sets its strictest bar for significance exactly there, while the more variable flanking sequence clears that bar more easily. Figure 2 G shows that per-motif seqlet counts agree closely with TF-MoDISco over the 26 shared motifs (Spearman *ρ* = 0.79), with points lying close to the diagonal rather than reallocating. On this benchmark the per-position null slightly over-identifies repressor motifs (14 against 3), the same tendency to be more permissive where signal is weak that drives the soybean promoter shift. We therefore interpret K562 as a positive control showing that CASCADE preserves established core-promoter motif recovery, rather than as evidence that it improves on TF-MoDISco.

Together, these results show that a position-specific null can redirect motif recovery when attribution background varies strongly across a sequence window while preserving established motif recovery in a well-characterized promoter-centered setting.

### 2.3 Applying CASCADE across cell-type-resolved attributions identifies broadly active, tissue-restricted and cell-type-restricted candidate sequence elements

Applying CASCADE independently to each of the 66 cell-type attribution tracks extends promoter-associated candidate sequence elements into a map of the cellular contexts in which candidate sequence elements influence model predictions. For each gene and cell type, CASCADE identifies short contiguous intervals whose attribution exceeds the cross-gene background at the same TSS-relative position. Intervals with centers within 10 bp are merged across cell types into cross-cell candidate sequence elements, and the resulting occupancy vector records the cell types in which each sequence element is detected.

Across 38,339 genes and 66 cell types, this procedure identifies approximately 1.39 million model-derived candidate sequence elements associated with 37,707 genes. Each candidate sequence element is specific to a single gene, since cell-type merging is performed only within that gene’s own attribution tracks; elements are never merged across genes, even when two genes carry similar or homologous sequence. Sequence elements detected in one or two cell types are classified as cell-type-restricted; sequence elements enriched within a single tissue are classified as tissue-restricted; elements detected across several tissues are classified as multi-tissue; and sequence elements present across most tissues are classified as broadly active or near-constitutive (Figure 3 D,E). These classes describe the specificity of model attribution rather than experimentally validated regulatory activity.

**Figure 3:**
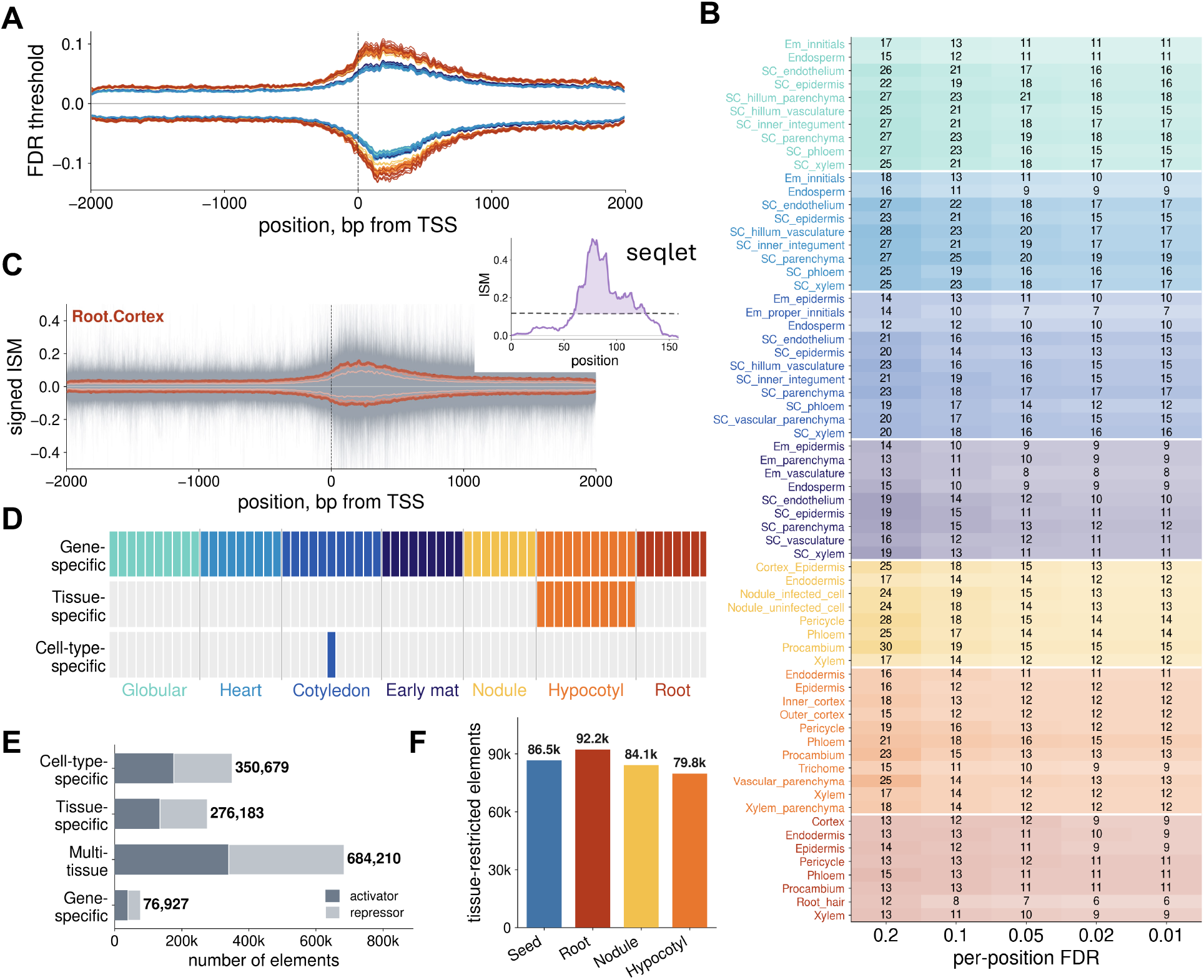
Position-specific significance thresholds resolve candidate regulatory elements across cell types, tissues and broadly active contexts. Running example, gene *Glyma*.*15G169300*. **A** Position-specific activator and repressor significance thresholds across the 66 cell types, colored by tissue and plotted relative to the TSS. **B** Number of seqlets per cell type across target FDR operating points, with rows grouped by tissue. **C** Signed in silico saturation mutagenesis tracks for sampled genes in the root cortex cell type, shown relative to position-specific significance thresholds at increasingly stringent operating points; a threshold crossing defines a candidate seqlet (inset). **D** Cellular-context classification. Cross-cell occupancy patterns define cell-type-restricted, tissue-restricted, multi-tissue and broadly active or near-constitutive candidate elements. **E** Candidate-element counts across the 38,339 TSS-centered gene windows, grouped by cellular-context class and predicted direction of mutational effect. **F** Number of tissue-restricted candidate elements identified within each organ group, with the four seed stages combined into a single seed organ.

Figure 1 E illustrates the resulting hierarchy using separate example loci. A broadly active candidate element produces a consistent predicted mutational effect across most cell types. A tissue-restricted element exceeds the position-specific background primarily within cell types belonging to one tissue, whereas a cell-type-restricted element is detected in only one or two cell types. Thus, when CASCADE is applied to cell-type-resolved attribution tracks, each candidate element is assigned a cellular-context profile rather than only a genomic position.

Resolving a single promoter across all 66 cell types reveals that neighboring candidate sequence elements can have distinct cellular-context profiles and predicted directions of effect. At the promoter of *Glyma*.*01G211000*, CASCADE identifies three nearby sequence elements with different occupancy patterns (Figure 4). One sequence element is detected across nearly all cell types, a second is restricted predominantly to early-nodule cell types, and a third is detected specifically in hypocotyl xylem (Figure 4 A). Expanding the same promoter along the cell-type axis shows that these sequence elements occupy distinct regions of the position-by-cell-type attribution surface (Figure 4 B,C). Their signed attribution scores further distinguish predicted activating from repressing effects. This example demonstrates how CASCADE separates spatially adjacent promoter sequences according to both their predicted direction of effect and the cellular contexts in which they influence model output.

**Figure 4:**
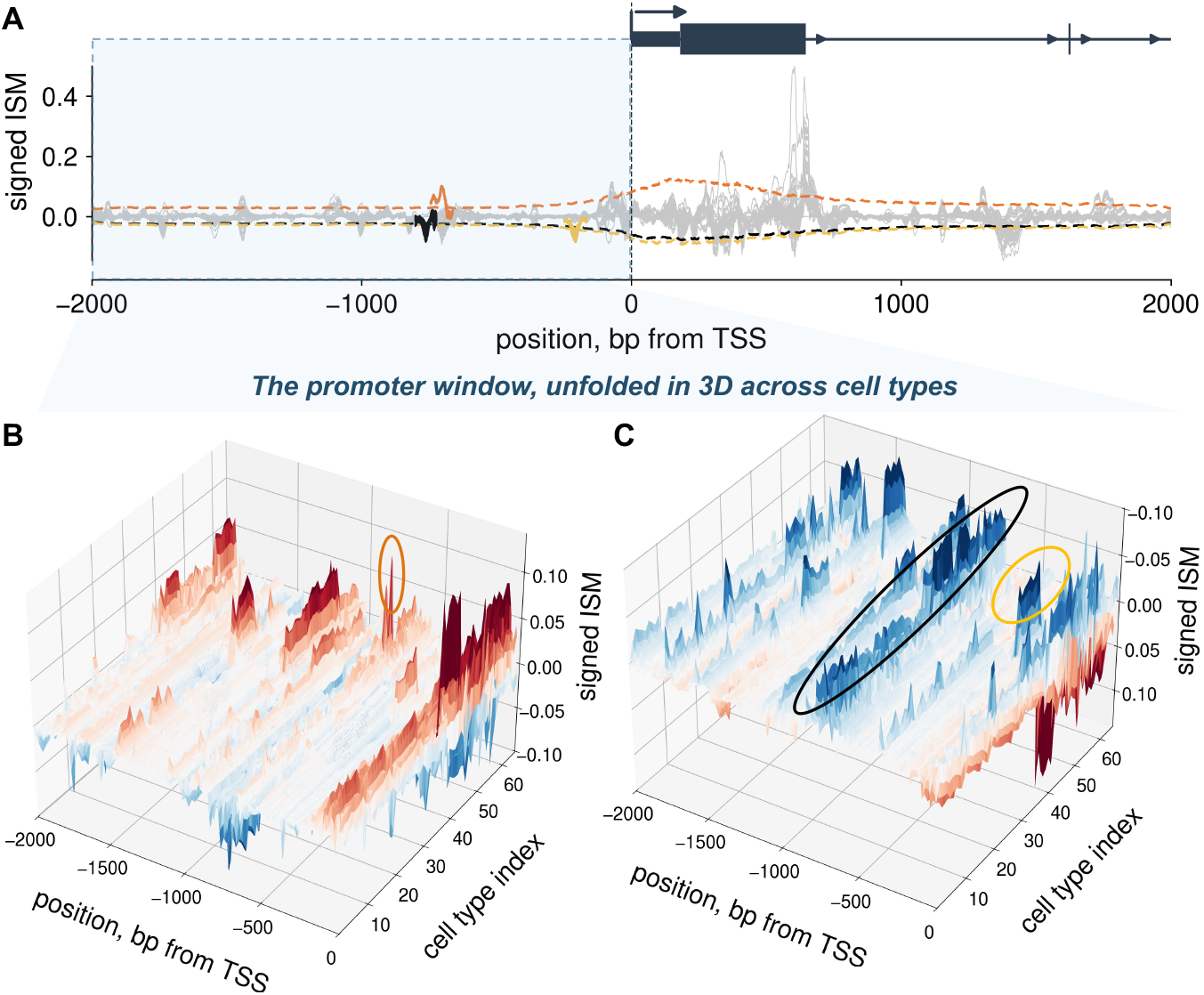
Neighboring candidate sequence elements within a single promoter exhibit distinct cellular-context profiles and predicted directions of effect. Running example, the promoter of *Glyma*.*01G211000*. **A** Signed *in silico* saturation mutagenesis tracks across all 66 cell types, with the gene model shown above. Three exemplar candidate elements are highlighted according to their cellular-context class: a broadly active or near-constitutive element in black, an early-nodule-restricted element in yellow and a hypocotyl-xylem-restricted element in orange. Each highlighted element exceeds its position-specific CASCADE significance threshold in a different set of cell types. The shaded promoter interval is expanded along the cell-type axis in **B** and **C. B**,**C** Position-by-cell-type surfaces of signed mutational effects across the promoter, with the 66 cell types ordered by tissue and developmental context. Activating predicted effects are shown above the surface in **B**, with the hypocotyl-xylem-restricted element indicated; repressing predicted effects are shown above the surface in **C**, with the early-nodule-restricted and broadly active elements indicated. The three elements influence model predictions in distinct cellular contexts despite occurring within the same promoter.

Tissue-restricted candidate ssquence elements are comparably abundant across the four soybean organs. Figure 3 F counts the tissue-restricted candidate elements identified within each organ group, with the four seed stages combined into a single seed organ. The organ totals are similar, and the developing seed carries a comparable number of tissue-restricted elements to root, nodule and hypocotyl. We report this organ-level total rather than a per-cell-type rate because the seed is the only organ sampled as a developmental time series, across four stages and 39 cell types, whereas each vegetative organ is sampled once across 8 to 11 cell types. A per-cell-type rate would divide a comparable organ total by many more cell types for the seed and would not reflect a difference in regulatory density across organs.

## 3 Discussion

CASCADE addresses a central limitation of attribution-based motif discovery by accounting for systematic variation in attribution magnitude around the transcription start site. Holding the underlying attributions, seqlet window, target operating point and downstream clustering fixed, replacing the pooled TF-MoDISco null with a position-specific cross-gene null redistributed motif instances from coding-proximal sequence toward the upstream promoter. The total number of seqlets remained similar between procedures, indicating that CASCADE primarily changes which sequence enters motif discovery rather than simply increasing the number of seqlets. This redistribution was strongest among method-exclusive motifs: 77% of CASCADE-exclusive motifs occurred in the promoter, compared with 12% of TF-MoDISco-exclusive motifs. At the same time, CASCADE preserved established core-promoter motif recovery on the human K562 ProCapNet benchmark. Applied independently across the 66 soybean cell-type attribution tracks, the framework identified approximately 1.39 million model-derived candidate sequence elements across 37,707 genes and resolved them into broadly active, multi-tissue, tissue-restricted and cell-type-restricted classes.

These results show that the statistical background used to identify seqlets shapes the regulatory grammar recovered from a fixed attribution map. Gene body regions contain strong and positionally constrained sequence signals, including translation initiation, untranslated-region, codonusage and transcript-processing features (Benegas et al., 2023; Agarwal and Shendure, 2020; Agarwal and Kelley, 2022; Hia et al., 2019; Peleke et al., 2024). A pooled background therefore favors regions with the largest absolute attribution effects, even when the biological question concerns promoter-associated regulation. CASCADE instead asks whether an attribution is unusual relative to other genes at the same TSS-relative position. This distinction allows lower-magnitude promoter signals to be evaluated against the appropriate local background rather than against the stronger attribution distribution found downstream of the TSS.

Recovering promoter-associated regulatory grammar is particularly important for plant genome editing because promoter edits can tune gene expression without altering the encoded protein. Editing *cis*-regulatory sequence can generate quantitative changes in expression and, in principle, modify when and where a gene is expressed rather than simply disrupting its function (RodrÍguez- Leal et al., 2017; Liu et al., 2021). A position-aware attribution framework can therefore help prioritize promoter intervals and sequence features for experimental editing, including candidates intended to alter expression in particular tissues or cell types. CASCADE does not establish that an individual edit will produce the predicted phenotype, but it narrows the sequence space to experimentally testable regulatory hypotheses.

The position-specific null is, by design, more permissive in regions where background attribution is lower, and part of the promoter shift therefore follows directly from the statistical formulation. CASCADE should consequently not be interpreted as universally superior to TF-MoDISco or as proving that every promoter-associated element is functional. Nevertheless, the CASCADE-exclusive motifs have information content comparable to the TF-MoDISco-exclusive motifs and match annotated transcription-factor families at least as frequently. The K562 analysis further shows that the position-specific procedure preserves established promoter motifs in a well-characterized benchmark, although it mildly increases repressor motifs and fragments the CTCF motif. Together, these observations support CASCADE as a position-aware alternative for questions focused specifically on promoter-associated grammar.

Applying CASCADE independently across cell-type-resolved attribution tracks converts promoter-associated sequences into cellular-context profiles. Each candidate sequence element is defined not only by its genomic position but also by the set of cell types in which its attribution exceeds the position-specific background. The resulting occupancy patterns distinguish regulatory elements detected broadly across the atlas from those restricted to a tissue or to one or two cell types. These classifications are operational descriptions of model attribution rather than direct measurements of regulatory activity: a broadly active element consistently influences model predictions across many cell types, whereas a cell-type-restricted element influences the model in only a limited cellular context. This additional axis transforms a static list of promoter-associated sequence intervals into a map of predicted regulatory specificity. Such cellular-context profiles could guide the selection of promoter edits intended to alter expression in a target tissue or cell type while minimizing effects in other contexts or provide biologically structured inputs into emerging deep learning architectures including biology-informed neural networks (BINNs) (Kontolati et al., 2025).

We do not interpret the organ-level element counts as evidence for a distinctive seed regulatory program. Because the developing seed is the only organ sampled across development, its lower per-cell-type sequence element density is confounded with the number of sampled cell states, and the comparable organ totals argue against a sparser seed program rather than for one. Distinguishing a genuinely more compact seed program from this sampling effect would require sampling the vegetative organs across matched developmental stages, which the current atlas does not provide.

The candidate sequence elements remain model-derived hypotheses rather than experimentally validated regulators. The analysis is restricted to a 4,000-bp window centered on each TSS and therefore excludes distal regulatory sequence and long-range chromatin interactions (Ricci et al., 2019; Lu et al., 2019; Rodgers-Melnick et al., 2016; Oka et al., 2017). Attribution scores were obtained from a single sequence-to-expression architecture and one reference sequence per gene, and attribution-based interpretations can vary across model architectures, random initializations and explanation methods (Novakovsky et al., 2023; Majdandzic et al., 2022; Toneyan et al., 2022; Adebayo et al., 2018). The activating and repressing labels describe the sign of the predicted expression response to mutation, not a demonstrated molecular mechanism. In addition, expression targets were cell-type-aggregated pseudobulk, where reads are aggregated by cell type into expression matrix profiles, and the mean per-cell-type, across-gene correlation does not by itself establish that the model fully captures within-gene expression differences among cell types. The cellular-context classes should therefore be interpreted as specificity in model attribution rather than definitive cell-type-specific regulatory activity.

Independent genomic data provide the most direct next test. The soybean atlas contains 303,199 cell-resolved accessible chromatin regions (Zhang et al., 2025), enabling evaluation of whether tissue- and cell-type-restricted CASCADE elements preferentially overlap chromatin accessibility in the corresponding cellular context. Single-cell chromatin accessibility atlases now available for maize, rice and other plants would further allow the cell-resolved regulatory grammar recovered here to be tested for conservation across species (Marand et al., 2021; Yan et al., 2024; Wang et al., 2025). Transcription-factor occupancy and natural regulatory variation would provide orthogonal evidence, whereas targeted CRISPR editing of prioritized promoter elements followed by cell- or tissue-resolved expression measurements would provide a direct functional test of the predicted regulatory effects. Replication across model architectures and training seeds would further establish whether the recovered elements and their cellular-context profiles are stable properties of the sequence rather than of a particular fitted model.

Natural allelic variation and multi-site perturbation provide complementary routes for extending CASCADE beyond single-reference, single-base interpretation. Applying the model to naturally occurring promoter haplotypes would produce multiple sequence contexts for each gene and support within-gene estimates of regulatory effects. Such allelic series would also enable direct prioritization of candidate *cis*-regulatory variants, a setting in which current sequence models can still predict the incorrect direction of effect (Huang et al., 2023). Multi-base perturbations could additionally reveal combinatorial motif interactions that single-base saturation mutagenesis cannot fully capture. Designed and natural-variant reporter libraries provide an experimental framework for evaluating both extensions (Rafi et al., 2025; Sample et al., 2019).

Together, these results establish CASCADE as a position-aware framework for extracting promoter-associated regulatory hypotheses from sequence-model attributions. The central methodological result is that the background used to identify seqlets determines which positional component of regulatory grammar enters motif discovery. Extending this comparison across cell-type-resolved model outputs produces a map of candidate promoter-associated elements and the cellular contexts in which they influence predicted expression. By prioritizing promoter sequences that can be experimentally perturbed without changing the encoded protein, this framework provides a path toward genome edits designed to tune the magnitude, timing and cellular specificity of gene expression in plants. The resulting atlas remains a set of model-derived hypotheses, but it focuses experimental validation and regulatory editing on defined sequence intervals with predicted context-specific effects.

## 4 Methods

### 4.1 Single-cell soybean atlas and expression

We used the single-cell RNA-seq atlas of *Glycine max* (Zhang et al., 2025). The atlas spans 66 cell types grouped into 7 tissues. Four are seed developmental stages, namely globular, heart, cotyledon, and early-maturation. The other three are non-seed tissues, namely nodule, hypocotyl, and root. Gene models, coordinates, and transcription start sites are taken from the Williams 82 Wm82.a4.v1 genome assembly and annotation (Phytozome 13, JGI, release 508) (Schmutz et al., 2010). It covers 38,339 protein-coding genes, the subset with a complete window around the transcription start site. Reads are aggregated by cell type to produce a pseudobulk expression matrix. We scale this matrix to counts-per-million and stabilize it by the transform log(1 + CPM). We refer to these log_1_*p*(CPM) values as expression throughout the paper.

### 4.2 Sequence backbone and per-base embedding

The backbone is GPN, a dilated-convolutional DNA language model trained by masked-language modeling without expression labels (Benegas et al., 2023). Starting from the Brassicales-pretrained checkpoint, we adapt it to soybean by continued masked-language-model training on *Glycine max* (Williams 82) coding, promoter, and 5^*′*^/3^*′*^ untranslated-region sequences, split into non-overlapping 512-bp windows. At each step 15% of nucleotides are replaced by a mask token and the model is trained to recover them by cross-entropy computed over the masked positions only. We optimize with AdamW (peak learning rate 5 *×* 10^*−*5^, weight decay 0.01, gradient norm clipped at 1.0, batch size 512 per GPU) under a cosine schedule with 5% linear warmup, in bf16 mixed precision on a 95*/*5 train/validation split, retaining the checkpoint with the lowest validation masked-token loss. The adapted backbone is then frozen. For gene *g* we take the window spanning TSS*±*2,000 and embed it at single-nucleotide resolution, giving a fixed feature matrix

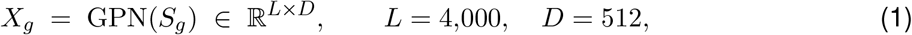

where *D* is the GPN hidden dimension and *S*_*g*_ is the reference sequence. The decoder is trained on the fixed embeddings.

### 4.3 Multi-task decoder

A convolutional decoder *f*_*θ*_ : ℝ^L*×*D^ *→* ℝ^C^ maps the gene embedding to predicted expression, one entry per cell type (*C* = 66), with internal width *d* = 768 and head width *h* = 1,024. With *X∈* ℝ^L*×*D^ transposed to channels-first, a two-layer convolutional encoder (kernels 3 and 5, each followed by batch normalization and ReLU, then dropout 0.3) is globally max-pooled over the length axis, layer-normalized, and linearly projected,

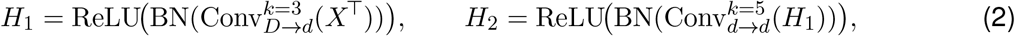

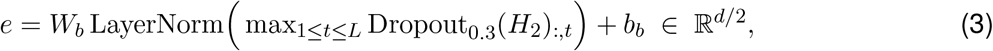

so that the global max-pool makes the representation invariant to where along the window a motif activation occurs. A linear low-rank bottleneck ***ℓ*** = *W*_**ℓ**_*e* + *b*_**ℓ**_ ∈ ℝ^d/4^ feeds a two-layer perceptron with a per-cell-type output row,

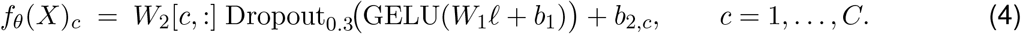

A single forward pass emits all *C* predictions. Cell identity enters only as the output index. Each cell type *c* corresponds to one row *W*_2_[*c*, :] of the final linear map, a per-cell readout of the same shared gene representation ***ℓ***. This multi-task readout is equivalent to conditioning a single-output model on a one-hot cell index at the readout, but costs one forward pass per gene rather than one per gene–cell pair, a *C*-fold reduction.

### 4.4 Training and cross-validation

The target for gene *g* is *Y*_*g*_ *∈* ℝ^C^ on the log_1_*p*(CPM) scale. The multi-task loss is the mean squared error over all gene–cell-type pairs in a batch *B* of genes,

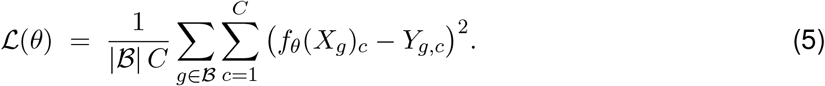

We optimize with AdamW (learning rate 5 *×* 10^*−*4^ on a constant schedule, weight decay 5*×* 10^*−*3^), an effective batch of 512 genes, up to 500 epochs with early stopping on validation MSE (patience 15), fixed seed 42, in single precision. We evaluate under 10-fold gene-level cross-validation, with the folds blocked by chromosome to guard against paralog and positional leakage between training and test. All *C* targets of a gene fall in the same partition, and held-out test genes are disjoint from training and validation. For each cell type we compute the Pearson correlation across held-out genes between *f*_*θ*_(*X*_*g*_)_*c*_ and *Y*_*g,c*_, then average across cell types (Figure 1b).

### 4.5 *In silico* saturation mutagenesis

We treat the trained model as a fixed function. For each of the *L* positions *i* and each alternative base *b*≠ref(*i*) we substitute the base, re-embed with the frozen backbone, and record the signed change in predicted expression,

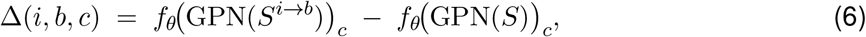

on the log_1_*p*(CPM) scale, giving a per-gene tensor in ℝ^L*×*4*×*C^ with the reference allele masked (Figure 1a). Two summaries are used downstream. For genome-wide element mapping we use the per-cell attribution

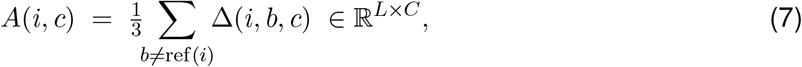

the mean over the three non-reference alleles, so that *A*(*i, c*) *>* 0 marks a repressing base (mutating it raises predicted expression) and *A*(*i, c*) *<* 0 an activating base. For display, the signed-in silico saturation mutagenesis tracks in the figures are drawn as *− A*, orienting them so that activating bases lie above zero and repressing bases below zero (Figure 1c). For motif discovery we use the cell-averaged, mean-centered hypothetical-contribution matrix in the TF-MoDISco format,

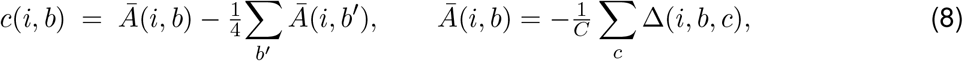

zeroed at padding positions; the negation makes *c* a contribution rather than a mutation effect. We compute these attributions for all 38,339 genes.

### 4.6 CASCADE: cross-gene, per-position FDR seqlet detection

Element and motif detection reuse the seqlet detector of TF-MoDISco (Shrikumar et al., 2018), the seqlet-detection paradigm introduced with base-resolution attribution models (Avsec et al., 2021b), changing *only* the null. For a cell *c* we stack *A*(·, *c*) over all *N* = 38,339 genes and form the length-*W* rolling-sum window score

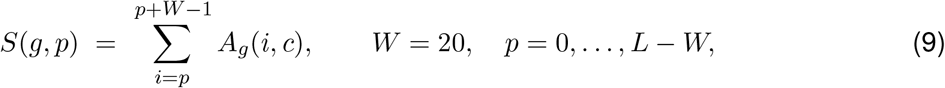

trimming a flank of *F* = 5 at the edges; the window center maps to genomic coordinate *p* + ***⌊*** *W/*2***⌋*** . TF-MoDISco fits one analytic null to the *pooled* set of window scores over all positions, giving two scalar thresholds. CASCADE keeps the identical estimator but fits it *per position*. At each column *p* it estimates a two-sided Laplace null 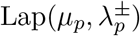 from 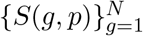by the mode-split, maximum-*λ* percentile fit of TF-MoDISco, Monte-Carlo sampling *M* = 3,000 null scores. Formally, TF-MoDISco assumes a position-homogeneous null (one (***µ***, *λ*^*±*^) for all *p*); CASCADE assumes a position-heterogeneous null (a separate 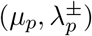 per position).

At target FDR *q* = 0.05, isotonic regression of the real-versus-null label on the score magnitude (the TF-MoDISco precision construction) yields, at each position, the smallest-magnitude score whose estimated precision reaches 1 *− q*,

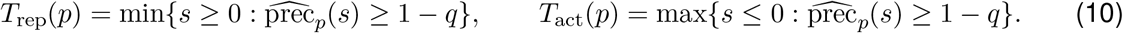

Positions with insufficient support return *±∞* and are linearly interpolated from finite neighbors; the threshold tracks are then smoothed with a length-*W* uniform filter to borrow strength across adjacent positions. A global pass-fraction constraint rescales the whole envelope by a single scalar *k* so that the overall fraction of passing windows lies in [0.03, 0.20], preserving the perposition shape. A window passes if *S*(*g, p*) ≥ *T*_rep_(*p*) or *S*(*g, p*) *≤T*_act_(*p*); seqlets are extracted per gene by greedy peak-picking on the masked | *S*| with non-maximum suppression of radius ***⌊****W/*2***⌋*** + *F* = 15, each a fixed *W* + 2*F* = 30-bp span signed by its central score. This yields a per-gene, per-cell set of significant seqlets.

### 4.7 Cross-cell elements and specificity classification

Running the detector for all *C* = 66 cell types gives, per gene and per sign, a set of seqlets. Seqlets whose centers lie within *T* = 10 bp are merged by single-linkage chaining into a cross-cell *element E*, a tolerance that keeps an element close to one binding footprint. With the tissue map *π* : *{*1, …, 66*} → {*1, …, 7*}* of sizes (*n*_*t*_) = (10, 9, 11, 9, 8, 11, 8) (globular, heart, cotyledon,early-maturation, nodule, hypocotyl, root;∑_*t*_ *n*_*t*_ = 66) an element has occupancy *o*(*E*) *∈ {*0, 1*}*^*66*^ number of occupied cell types *b*(*E*) =∑_c_ *o*_*c*_ (*E*), per-tissue counts 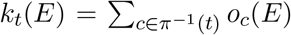, and within-tissue rates *r*_*t*_(*E*) = *k*_*t*_(*E*)*/n*_*t*_. Tissue enrichment is tested against a hypergeometric null that models the *b*(*E*) occupied cell types as draws without replacement from the 66, so that *k*_*t*_(*E*) *∼* Hypergeom(66, *n*_*t*_, *b*(*E*)) with tail probability *p*_*t*_(*E*) = Pr(*X≥ k*_*t*_(*E*)), element statistic *p*_min_(*E*) = min_*t*_ *p*_*t*_(*E*) at *t*^**⋆**^(*E*) = arg min_*t*_ *p*_*t*_(*E*), and Benjamini–Hochberg value *q*(*E*) over all elements with *b ≥* 3. With *q*^**⋆**^ = 0.05, *ϕ* = 0.5, ***Ψ*** = 0.2, *γ* = 0.3, *m* = 6, each element is classified per sign as

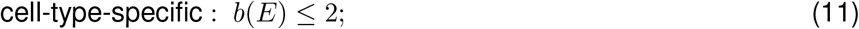

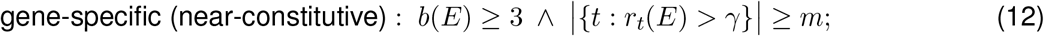

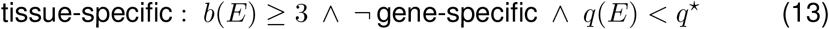

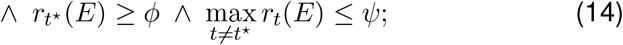

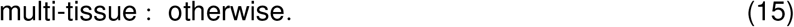

These are operational strata over the model’s cross-cell attributions. Occupancy is a model-internal detection, and this count is not normalized for per-cell detection power (the hypergeometric test corrects for tissue size *n*_*t*_ but not for tissue-correlated detectability). Specificity here is therefore specificity of model attribution, tested against a null only for the tissue-specific class.

### 4.8 Comparison of seqlet-detection nulls and the ProCapNet benchmark

To isolate the effect of the null on motif discovery we ran two motif-discovery passes that differ only in the detection null. The baseline is TF-MoDISco with its pooled, position-flat FDR null. The variant, CASCADE, replaces that null with the per-position FDR null described above; the window, flank, target FDR and the downstream seqlet clustering are held identical, so any difference in the recovered motifs is attributable to the null alone. For soybean we ran both passes on the cell-averaged in silico saturation mutagenesis attributions, pooled over the union of the 10 cross-validation folds and all 38,339 genes. We matched motifs one-to-one across passes by contribution-weight-matrix correlation, designating a pair shared at a correlation of at least 0.80, and screened the method-exclusive motifs by median seqlet position relative to the TSS, per-position information content, low-complexity content, and TOMTOM matches to annotated transcription-factor families. Per-position information content is the Schneider–Stephens sequence-logo information (Schneider and Stephens, 1990), 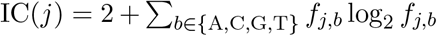 bits, where *f*_*j,·*_ is the position frequency matrix of the motif’s aligned seqlets restricted to its information-content-trimmed core. We report the mean over core positions (Figure 2d), and exclude low-complexity motifs, those dominated by a single base, from this comparison. As an external, base-resolution positive control we repeated the comparison on the deposited human K562 PRO-cap model of ProCapNet (Cochran et al., 2024), whose core-promoter transcription factors are annotated, and scored recovery as the best-match contribution-weight-matrix correlation of each pass to the reference motifs.

## Acknowledgements

We would like to thank Drs. Jie Yao, Xuan Zhang, and Hongwoo Lee for feedback on this study.

R.J.S. is supported by the National Science Foundation (IOS-1856627) and the Office of Research at the University of Georgia. We also thank the Georgia Advanced Computing Resource Center (GACRC) at the University of Georgia for computational resources.

## 4.9 Data and code availability

All code and links to larger datasets, such as the single-cell soybean atlas and per-gene attribution library can be found via the GitHub repository: CompAgLab/CASCADE

## Supplementary Information

We assessed the sequence-to-expression model as a high-versus-low expression classifier, following the DeepCRE protocol (Peleke et al., 2024). For each tissue, we defined expression as the mean log_1_*p*(CPM) across that tissue’s single-cell types. Genes in the bottom expression quartile were labeled low and genes in the top quartile high, and the middle 50% were discarded, giving two balanced categories per tissue. We then ranked genes by the model’s held-out out-of-fold predicted expression and scored the separation of the high and low categories by the area under the receiver-operating-characteristic curve, without training any additional classifier.

**Figure S1:**
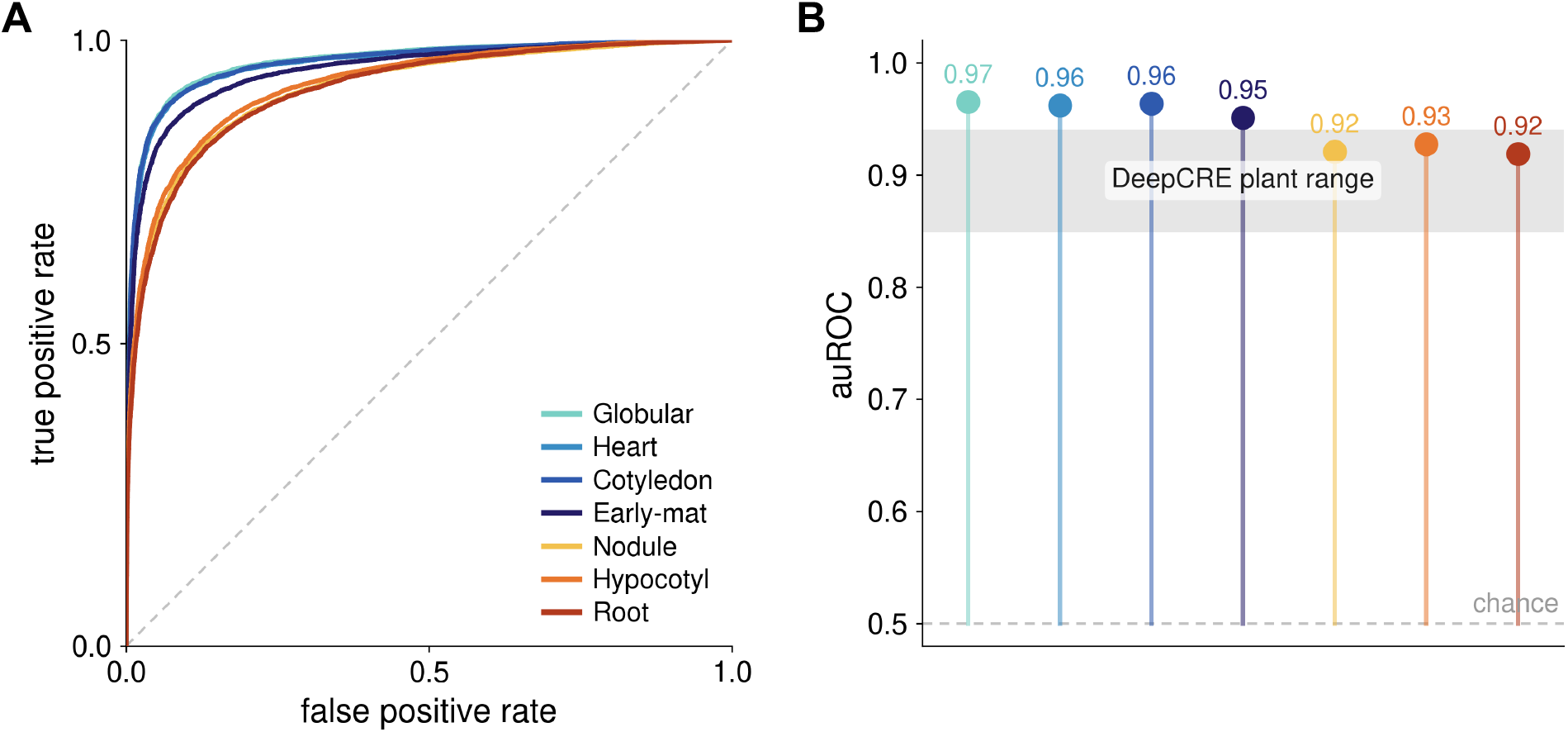
The sequence-to-expression model separates high- from low-expressed genes across all tissues. (A) Per-tissue receiver-operating-characteristic curves. (B) Per-tissue area under the curve, with the shaded band marking the reported plant range and the dashed line marking chance.

